# An mTOR Inhibitor Discovery System Using the Growth of Drug-Sensitized Yeast Strains

**DOI:** 10.1101/2025.01.03.631268

**Authors:** Anna K. Breen, Sarah Thomas, David Beckett, Matthew Agsalud, Graham Gingras, Judd Williams, Brian M. Wasko

**Affiliations:** Department of Biomedical Sciences, Western University of Health Sciences, Lebanon, OR 97355

## Abstract

Inhibition of the target of rapamycin (TOR/mTOR) protein kinase by the drug rapamycin extends lifespan and healthspan across diverse species. However, rapamycin has potential off-target and side effects that warrant the discovery of additional TOR inhibitors. TOR was initially discovered in *Saccharomyces cerevisiae* (yeast) which contains two TOR paralogs, *TOR1* and *TOR2*. Yeast lacking functional Tor1 are viable but are hypersensitive to growth inhibition by TORC1 inhibitors, which is a property of yeast that can be exploited to identify TOR inhibitors. Additionally, yeast lacking FK506-sensitive proline rotamase (*FPR1*) or containing a *tor1-1* allele (a mutation in the Fpr1-rapamycin binding domain of Tor1) are robustly and selectively resistant to rapamycin and analogs that allosterically inhibit TOR activity via an *FPR1*-dependent mechanism.

To facilitate the identification of TOR inhibitors, we generated a panel of yeast strains with mutations in TOR pathway genes combined with the removal of 12 additional genes involved in drug efflux. This creates a drug sensitive strain background that can sensitively and effectively identify TOR inhibitors.

In a wildtype yeast strain background, 25 µM of Torin1 and 100 µM of GSK2126458 (omipalisib) are necessary to observe *TOR1*-dependent growth inhibition by these known TOR inhibitors. In contrast, 100 nM Torin1 and 500 nM GSK2126458 (omipalisib) are sufficient to identify *TOR1-*dependent growth inhibition in the drug sensitized background. This represents a 200-fold and 250-fold increase in detection sensitivity for Torin1 and GSK2126458, respectively. Additionally, for the TOR inhibitor AZD8055, the drug sensitive system resolves that the compound results in *TOR1-*dependent growth sensitivity at 100 µM, whereas no growth inhibition is observed in a wildtype yeast strain background. Our platform also identifies the caffeine analog aminophylline as a *TOR1*-dependent growth inhibitor via selective *tor1* growth sensitivity. We also tested nebivolol, isoliquiritigenin, canagliflozin, withaferin A, ganoderic acid A, and taurine, and found no evidence for TOR inhibition using our yeast growth-based model.

Our results demonstrate that this system is highly effective at identifying compounds that inhibit the TOR pathway. It offers a rapid, cost-efficient, and sensitive tool for drug discovery, with the potential to expedite the identification of new TOR inhibitors that could serve as geroprotective and/or anti-cancer agents.

## Introduction

The serine/threonine protein kinase known as the target of rapamycin (TOR or mechanistic/mammalian mTOR) was originally named for its discovery in *Saccharomyces cerevisiae* (yeast), where mutations in the two yeast TOR genes were found to suppress rapamycin-mediated inhibition of growth (Heitman et al. 1991; Rivera and Heitman 2023). mTOR is a highly conserved protein kinase that is a master regulator of cell growth and proliferation that responds to multiple external stimuli.

Genetic reduction of the mTOR signaling pathway extends the lifespan of *Caenorhabditis elegans* (Vellai et al. 2003), *Drosophila melanogaste*r (Kapahi et al. 2004), yeast (Kaeberlein et al. 2005; Powers et al. 2006), and mice (Wu et al. 2013). Pharmacological inhibition of TOR by rapamycin extends lifespan in yeast (Powers et al. 2006), flies (Bjedov et al. 2010), and mice (Harrison et al. 2009). Interestingly, even a short duration of rapamycin treatment in middle-aged mice is sufficient to extend longevity (Bitto et al. 2016).

The mTOR kinase functions within two distinct protein complexes, TORC1 and TORC2. TORC1 is a nutrient responsive kinase that promotes anabolic (e.g., protein synthesis) and regulates catabolic (e.g., autophagy) processes to promote cell growth in response to appropriate nutrients(Liu and Sabatini 2020). TORC2 is less well studied and has distinct downstream effects (Gaubitz et al. 2016). Given that mTOR is a master regulator of cell growth and cell division, rapamycin and other TOR inhibitors are also of interest for cancer treatment (Popova and Jücker 2021) and prevention (Blagosklonny 2023).

Rapamycin and its analogs (rapalogs) show promise as geroprotective and anti-cancer therapeutics, but multiple limitations may temper their application. One common concern is that rapamycin was first FDA-approved for the prophylaxis of organ transplant rejection, and thus its immunosuppressive properties may increase susceptibility to infections. Despite this concern, some accumulating evidence suggests that rapamycin may have immunomodulatory benefits in the aged population. A meta-analysis of mouse studies concluded that treatment with rapamycin significantly increases the short-term survival of mice in response to acute pathogen infection (Phillips and Simons 2023). In addition, a recent systematic review of human studies supports that rapamycin or rapalog treatment can promote beneficial effects for the aging immune system (Lee et al. 2024). In a survey study of off-label use of rapamycin (consisting primarily of once weekly dosing), there was a statistically significant increase in the incidence of mouth ulcers reported by rapamycin users compared to a control group, as well as a non-statistically significant trend toward higher infection rates (Kaeberlein et al. 2023). These studies highlight that the FDA-approval of rapamycin as an immunosuppressant is not sufficient to dismiss its use as geroprotective agent, though concerns persist.

Another concern is that chronic rapamycin treatment can result in metabolic disturbances due to off-target effects. This off-target effect is attributed to rapamycin binding free mTOR and preventing its assembly into TORC2 (Sarbassov et al. 2006), thereby reducing TORC2 activity and impairing glucose tolerance (the ability to regulate blood sugar levels after a glucose load) in mice (Lamming et al. 2012). Modifying the treatment regimen of rapamycin can mitigate these adverse effects in animal models, improving both glucose regulation and immune function (Arriola Apelo et al. 2016). In humans, intermittent dosing similarly reduces rapamycin-induced glucose intolerance, although it does not fully eliminate it (Baghdadi et al. 2024).

In order to accelerate the identification of novel mTOR inhibitors, the development of cost-effective and rapid assays are warranted. Ideal assays should be sensitive, specific, and amenable to high-throughput screening of molecular libraries. Developing novel processes for identifying TOR inhibitors can expedite drug discovery and reveal promising candidates for further investigation as potential geroprotective agents. Systems that can evaluate high-throughput combinations of compounds will also be useful.

*S. cerevisiae* contains two TOR genes, *TOR1* and *TOR2* that encode proteins which primarily function within the TORC1 and TORC2 complexes, respectively. Yeast lacking *TOR1* are viable but are hypersensitive to TORC1 inhibitors such as rapamycin due to a low residual activity of the Tor2 protein within the TORC1 complex (Figure 1A). This property of yeast can be exploited to identify TORC1 inhibitors by assessing for compounds that selectively inhibit growth of a yeast strain lacking a functional Tor1 protein (Lee et al. 2017). Compounds that do not inhibit TORC1 activity are expected to similarly inhibit yeast growth in both Tor1 deficient and control strains or to have no growth inhibitory properties at all.

**Figure 1.**
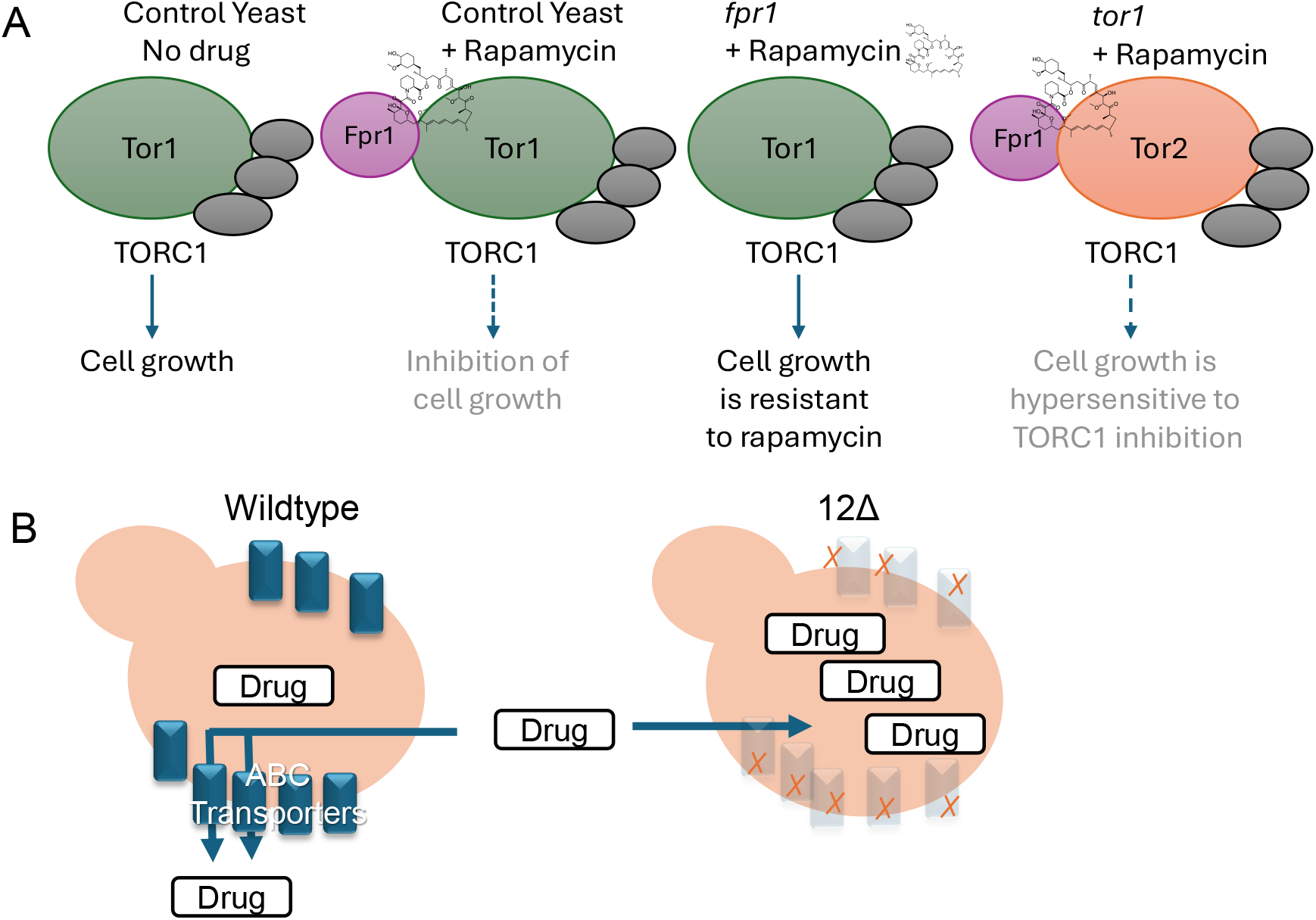
Representation of yeast strains used in this study. **A)** Growth inhibition by rapamycin and rapalogs depends on Fpr1 for the allosteric inhibition of TOR complex 1 (TORC1). Wildtype yeast are sensitive to rapamycin-mediated growth inhibition, while *fpr1*-deficient mutant yeast exhibit robust resistance to rapamycin and rapalogs due to the requirement of Fpr1 for the allosteric drug mechanism involving binding with rapamycin and the Tor protein. In contrast, *tor1*-deficient mutant yeast are hypersensitive to rapamycin and other TORC1 inhibitors. Viability of yeast *tor1* mutants and the hypersensitivity to TORC1 inhibitors arises due to low residual activity of the yeast Tor2 protein functioning within TORC1, which remains susceptible to inhibition. Grey circles represent other protein components of the TORC1 complex. **B)** Schematic comparing the wild-type strain background (BY4742 or BY4741) to the drug-sensitized 12Δ strain, which lacks multiple ABC transporters that can function as drug efflux pumps.

Additionally, yeast lacking functional Fpr1 or containing a *tor1-1* allele (a S1972R mutation in the Fpr1 binding domain of Tor1) are robustly and selectively resistant to rapamycin. This resistance arises because rapamycin allosterically inhibits TORC1 via an Fpr1 (FKBP12) dependent mechanism (Heitman et al. 1991). Of note, the interaction of Fpr1 with TORC1 is primarily a drug-dependent phenomenon and Fpr1 has no known endogenous role in TORC1 regulation in the absence of rapamycin.

As a unicellular organism evolved to survive in complex and harsh environments, yeast possess robust defensive mechanisms that help afford protection from their surroundings. These systems include numerous ATP-binding cassette (ABC) transport pumps capable of exporting potential harmful substances including many drugs. Chinen et al. have developed a strain of yeast lacking 12 genes involved in drug efflux, which display increased sensitivity to a variety of compounds (Chinen et al. 2014). Using the 12geneΔhsr strain (referred to as 12Δ within this manuscript, Figure 1B), the TOR pathway was modified to create a drug sensitive system that can identify agents that inhibit yeast growth in a TORC1 dependent manner.

We sought to generate a drug-sensitized yeast model for use in the identification of TOR inhibitors. We hypothesized that using a drug efflux deficient strain containing TOR pathway associated mutations would allow for a system that will maintain selectivity while increasing sensitivity to growth inhibition via TOR inhibitors. Using multiple compounds known to inhibit the TOR pathway, we provide a proof of principle that the developed system can functionally be used to increase sensitivity for the identification of TOR inhibitors. We also tested a panel of molecules previously implicated to impact mTOR and/or lifespan, including: nebivolol, isoliquiritigenin, α-lipoic acid, canagliflozin, withaferin A, ganoderic acid A, and taurine; and found no supportive evidence for TOR inhibition using our yeast growth based assay.

## Results

Rapamycin and its analogs result in hypersensitivity with *tor1* mutants, while *fpr1* and *tor1-1* mutants exhibit robust resistance.

A *tor1* strain is hypersensitive to rapamycin and the analogs ridaforolimus, temsirolimus, and everolimus, while *fpr1* or *tor1-1* mutant strains are strongly resistant (Heitman et al. 1991; Lee et al. 2017). We set out to determine if the same profile of growth is retained in the 12Δ drug sensitive genetic background. Growth curves for strains in the absence of drugs are shown for wildtype (Figure 2A) and 12Δ yeast strains (Figure 2B). Rapamycin strongly inhibited growth of wildtype (Figure 2C) and the 12Δ strain (Figure 2D) at 5 and 20 nM and the *tor1* deficient strains displayed an increased sensitivity to rapamycin at these concentrations, while the *fpr1* and *tor1-1* strains exhibited robust resistance to rapamycin.

**Figure 2.**
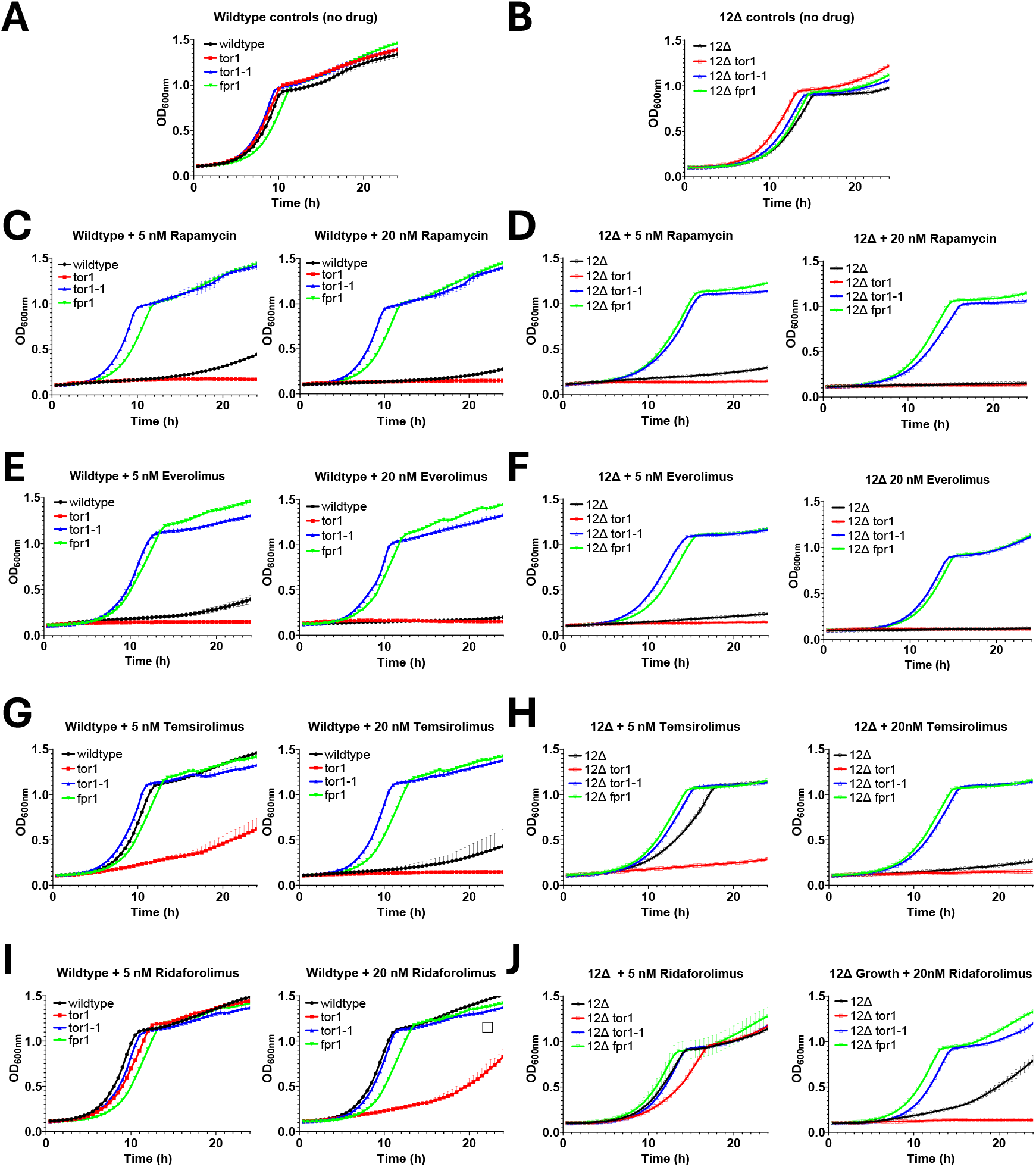
Rapamycin and its analogs result in hypersensitivity in *tor1* deficient mutants, while *fpr1* and *tor1-1* mutants exhibit robust resistance. Growth curves are shown for **A)** wildtype BY4742 haploids without drug, **B)** 12Δ yeast without drug, **C)** wildtype treated with 5 nM or 20 nM rapamycin, **D)** 12Δ treated with 5 nM or 20 nM rapamycin, **E)** wildtype treated with 5 nM or 20 nM everolimus, **F)** 12Δ treated with 5 nM or 20 nM everolimus, **G)** wildtype BY4742 treated with 5 nM or 20 nM temsirolimus, **H)** 12Δ treated with 5 nM or 20 nM temsirolimus, **I)** wildtype BY4742 treated with 5 nM or 20 nM ridaforolimus, and **J)** 12Δ treated with 5 nM or 20 nM ridaforolimus. OD600 is the optical density measured at 600 nM. Closed shapes indicate wildtype strains and open shapes represent 12Δ strains. Black circles are control strains, red squares are *tor1* disruption mutant strains, blue triangles are *tor1-1* (Tor1-S1972R) mutants, and green triangles are *fpr1* disruption mutants. Error bars represent standard deviation.

Everolimus strongly inhibited growth of wildtype (Figure 2E) and the 12Δ strain (Figure 2F) at 5 and 20 nM, and the *tor1* deficient strains also displayed an increased sensitivity to everolimus at these concentrations, while the *fpr1* and *tor1-1* strains exhibited strong resistance to everolimus.

Temsirolimus was less potent compared to rapamycin and everolimus at 5 nM in both wildtype (Figure 2G) and 12Δ backgrounds (Figure 2H), while an increased sensitivity to growth inhibition was observable in the *tor1* deficient strains. The *fpr1* and *tor1-1* strains exhibited resistance to growth inhibition in both the wildtype and 12Δ background.

Ridaforolimus appeared less potent than rapamycin and other rapalogs at inhibiting the growth of control strains in both wildtype (Figure 2I) and the 12Δ background (Figure 2J), with an increased sensitivity observed in the 12Δ compared to control background at 20 nM. The *tor1* strain exhibited enhanced sensitivity compared to the control strains in both the wildtype and 12Δ backgrounds at 20 nM ridaforolimus. The *fpr1* and *tor1-1* strains exhibited marked resistance to ridaforolimus in both wildtype and 12Δ.

The 12Δ strain background strongly increases sensitivity to ATP competitive TOR inhibitors: AZD8055, Torin1, and GSK2126458 (omipalisib).

ATP competitive inhibitors constitute another class of mTOR inhibitors distinct from rapamycin and rapalogs. AZD8055 (Chresta et al. 2010) and Torin1 (Liu et al. 2010) are ATP-competitive TORC1/TORC2 inhibitors, as is GSK2126458 (omipalisib), which additionally inhibits phosphoinositide 3-kinase (PI3K) (Knight et al. 2010). Growth curves for no drug controls are indicated in Figure 3A, and the compound AZD8055 did not inhibit growth of the control or *tor1* strain in the wildtype background at 100 µM (Figure 3B). In the 12Δ yeast, 100 µM of AZD8055 inhibited growth of the control strain and preferential inhibition in the *tor1* strain was observed. At 25 µM, Torin1 had a growth inhibitory effect in the control wildtype strain and a selective inhibition of the *tor1* strain was observed, while no growth of either the control or *tor1* strain was observed at 25 µM torin1 for the 12Δ strains (Figure 3C). Treatment with 100 nM Torin1 resulted in no growth inhibition observed in the wildtype strain background, but the 12Δ control strain was inhibited and a selective inhibition of the 12Δ *tor1* strain was observable (Figure 3D). Treatment with 50 µM GSK2126458 resulted in a preferential inhibition of the *tor1* strain compared to the control strain in the wildtype background, and no growth was observed in the 12Δ control or *tor1* strains (Figure 3E). At 500 nM, GSK2126458 did not inhibit growth of the wildtype background strains, but inhibited the growth of the 12Δ strains, with a preferential inhibition of the 12Δ *tor1* strain observed (Figure 3F). All of the tested ATP competitive inhibitors were identified as having *tor1*-dependent sensitivities using the drug sensitized system, while a wildtype system was unable to identify one even at high concentration. The two others were detected as having *tor1* sensitive growth at ∼200 fold lower concentrations in the 12Δ strain.

**Figure 3.**
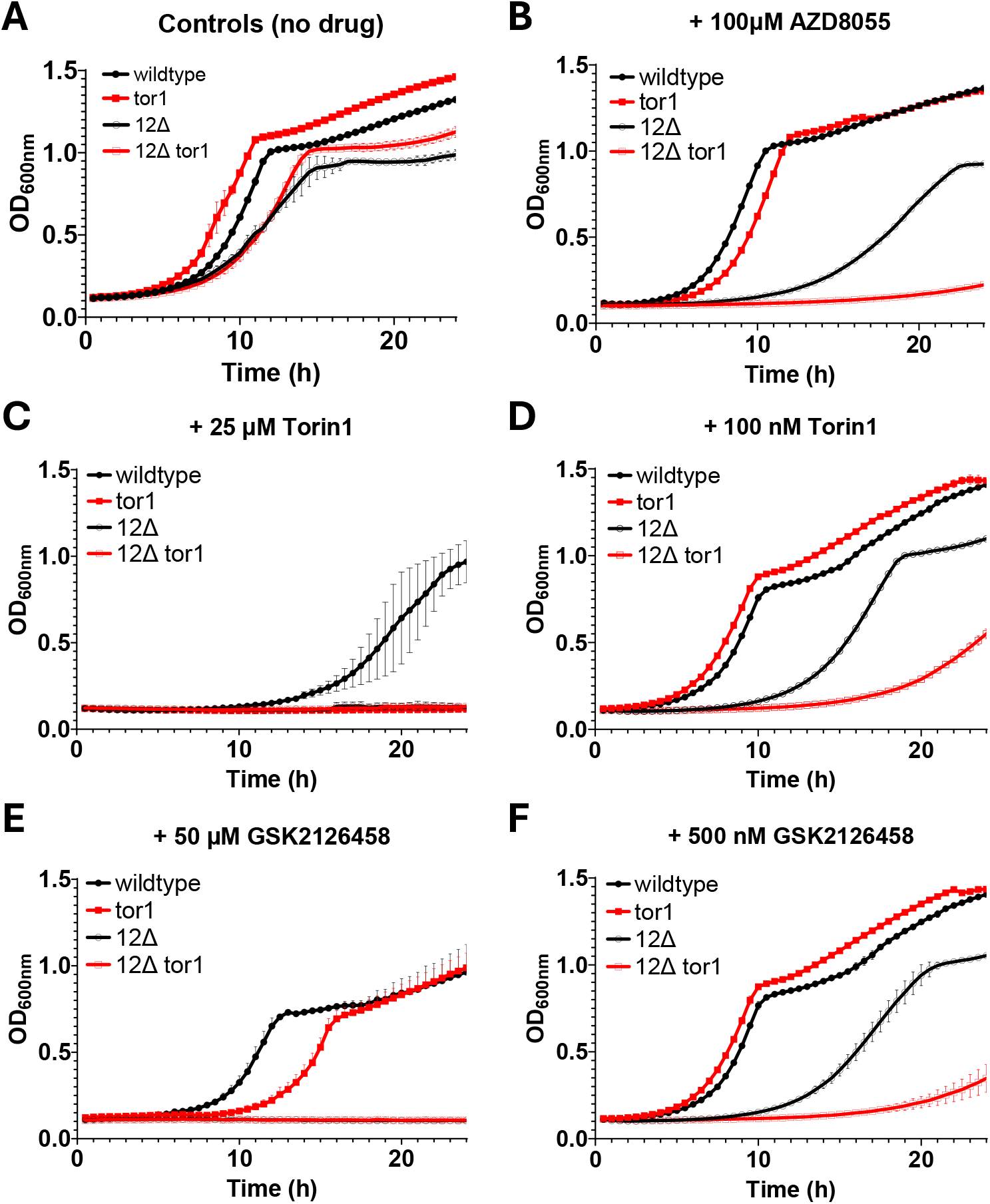
The ATP competitive TOR inhibitors AZD8055, Torin1, and GSK2126458 (omipalisib) are more potently identifiable as *tor1*-dependent growth inhibitors in the 12Δ background. Growth curves on YPD media for wildtype BY4742 and 12Δ treated with **A)** no drug, **B)** 100 µM AZD8055, **C)** 25 µM Torin1, **D)** 100 nM Torin1, **E)** 50 µM GSK2126458 (omipalisib), and **F)** 500 nM GSK2126458 (omipalisib). OD600 is the optical density measured at 600 nM. Filled shapes are wildtype BY4742 strains and open shapes are 12Δ strains. Black circles represent control strains and red squares represent *tor1* disruption mutants. Error bars represent standard deviation.

The 12Δ strain displays increased growth sensitivity to caffeine and its analog aminophylline and *tor1* mutants are selectively sensitive to growth inhibition.

Caffeine has previously been found to inhibit yeast growth in a *tor1*-dependent manner (Reinke et al. 2006). Controls without drug are indicated for the wildtype (Figure 4A) and 12Δ strain backgrounds (Figure 4B) with control, *tor1*, and an additional strain containing a *TOR1* allele encoding for a I1954V amino acid substitution previously identified as a caffeine resistant mutation (Reinke et al. 2006). Caffeine at 10 mM inhibited growth of the control strain of a wildtype background, while the *tor1* deficient mutant was preferentially inhibited, and the I1954V mutation conferred resistance (Figure 4C). In the 12Δ background with 10 mM caffeine, an increased sensitivity of the 12Δ control strain was observed compared to the wildtype strain, and the *tor1* deficient mutant also displayed a potent growth inhibition and the I1954V allele retained resistance (Figure 4D). The analog of caffeine, aminophylline, at 10 mM displayed moderate preferential inhibition of growth of the *tor1* deficient strain in the wildtype background, however no resistance was noted when Tor1 contained the I1954V mutation (Figure 4E). In the 12Δ background, 10mM aminophylline inhibited growth of the control strain and Tor1-I1954V mutant strain, and an increased sensitivity of the *tor1* deficient mutant was observable (Figure 4F). No resistance was observed for caffeine or aminophylline in *fpr1* and *tor1-1* strains (data not shown).

**Figure 4.**
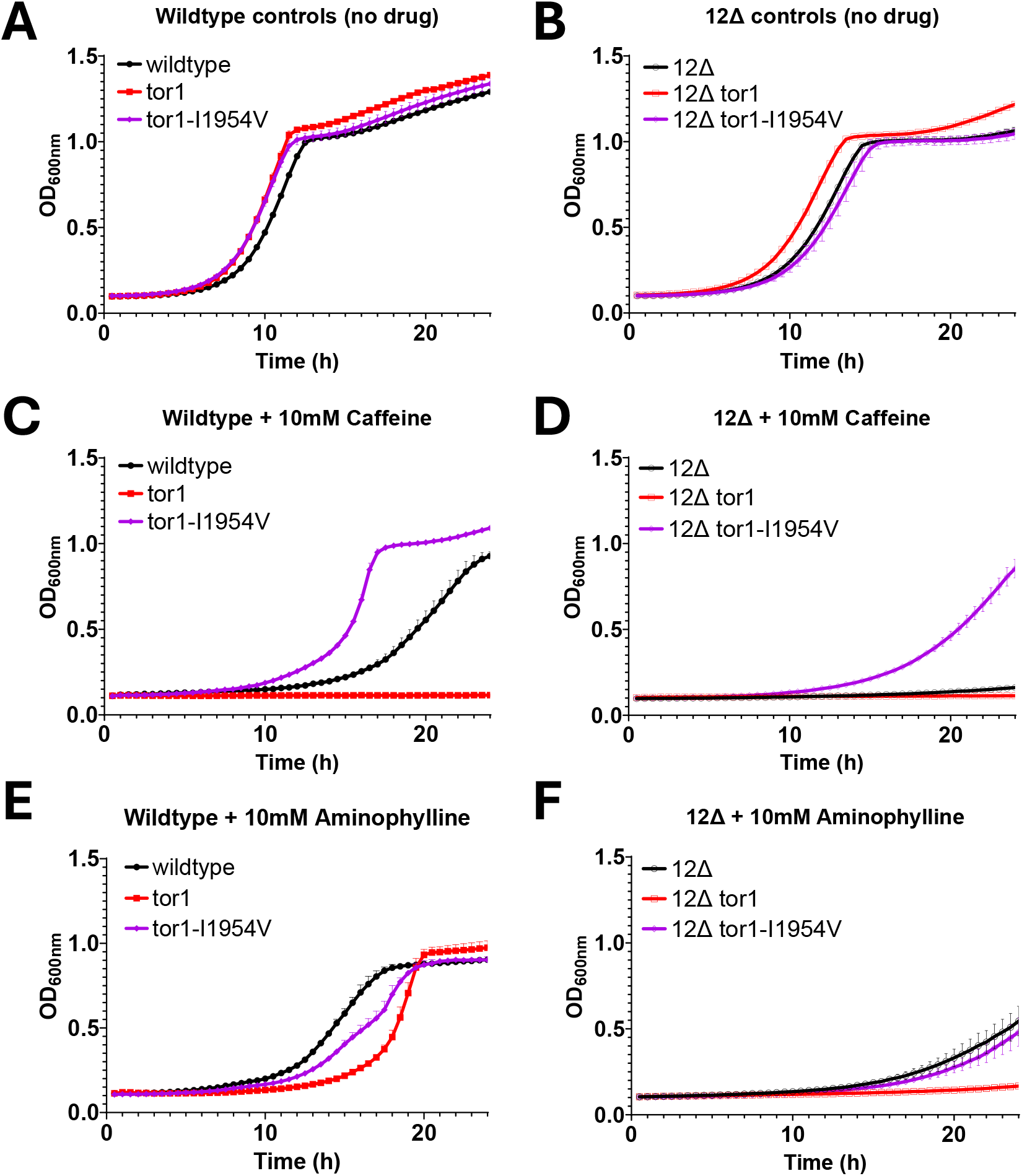
The 12Δ strain displays increased growth sensitivity to caffeine and its analog aminophylline, and *tor1* mutants are selectively sensitive to growth inhibition. Growth curves on YPD media for **A)** wildtype control BY4742, *tor1*, and *tor1-2* (Tor1-I1954V) without drug, **B)** 12Δ control, *tor1*, and *tor1-2* (Tor1-I1954V) without drug, **C)** wildtype BY4742, *tor1*, and *tor1-2* (Tor1-I1954V) with 10 mM caffeine, **D)** 12Δ control, *tor1*, and *tor1-2* (Tor1-I1954V) with 10 mM caffeine, **E)** wildtype control BY4742, *tor1*, and Tor1-I1954V with 10 mM aminophylline, and **F)** 12Δ control, *tor1*, and Tor1-I1954V with 10 mM aminophylline. OD600 is the optical density measured at 600 nM. Filled shapes are wildtype BY4742 strains and open shapes are 12Δ strains. Black circle shapes are control strains, red squares are *tor1* strains, and purple diamonds are tor1-I1954V mutants. Error bars represent standard deviation.

We assessed a variety of additional compounds with potential TORC1 effects using our drug sensitive model (Supplemental Table 1). We found that ganoderic acid A, α-lipoic acid, and taurine did not affect growth of the 12Δ strain at the concentrations tested. Nebivolol, Canagiflozin, Isoliqisoliquiritigenin, and Withaferin A were found to selectively inhibit growth in the 12Δ background, however no observable selective sensitivity in the *tor1* strain was observed at the tested concentrations (Supplemental Table 1).

## Discussion

Rapamycin and rapalogs function as allosteric mTOR inhibitors that require binding with the protein FKBP12/Fpr1 and inhibit mTOR by binding to the Fkbp12 binding domain (FBD) within TOR. In the absence of Fpr1, or when a *tor1-1* (S1972R) mutation is present, yeast cells become insensitive to rapamycin up to ∼ 100 µM (Rivera and Heitman 2023). With rapamycin and other rapalogs, the 12Δ background strains retained relatively similar growth profiles overall to the control ABC transporter proficient strain. This suggests that rapamycin and analogs are not likely to be efficiently effluxed from yeast cells, perhaps in part due to the tight binding with Fpr1, which may contribute to the relative potency of these molecules.

The 2nd generation mTOR inhibitors function as ATP competitive inhibitors of TOR with variable specificity. AZD8055 (Chresta et al. 2010) and Torin1 (Liu et al. 2010) are ATP-competitive TORC1/TORC2 inhibitors. GSK2126458 (omipalisib) is an ATP-competitive inhibitor of PI3K and mTORC1/mTORC2 (Knight et al. 2010). The increased sensitivity of yeast lacking multiple plasma membrane ABC-transporter pumps to these ATP-dependent mTOR inhibitors suggests that these compounds are normally actively effluxed from yeast cells. GSK2126458 at 10 µM is reported to extend the lifespan of *C. elegans* when treatment is started at adulthood (Phelps et al. 2024), and it is also found to extend the lifespan of *C. elegans* by 35% by the ongoing Million Molecule Challenge (Biomedical 2024), which uses a wormbot lifespan assay (Pitt et al. 2019) for high-throughput longevity drug discovery. Of note, rapamycin effects on *C. elegans* longevity are potentially confounded by precipitation issues at the reported concentrations (Lucanic et al. 2017; Phelps et al. 2024). Whether the GSK2126458 lifespan benefit is fully or partially dependent on a non-mTORC1 target such as PI3K may be of continued interest. Particularly of note, the first identified genetic longevity modifier in *C. elegans*, age-1, is a PI3K (Friedman and Johnson 1988; Morris et al. 1996). Caffeine and its analogs also extend worm longevity and can dually inhibit mTOR and PI3K (discussed further below).

Caffeine is a widely consumed substance and a large body of literature surrounds the numerous biological effects. In yeast, we find that both caffeine and the analog aminophylline inhibit yeast growth in a *tor1*-dependent manner and the 12Δ strain is sensitized to both. Loss of *TOR1* has previously been found to sensitize yeast cells to caffeine (Reinke et al. 2006; Lee et al. 2017), and growth suppression and adaptation experiments using high concentrations of caffeine have identified mutations in *TOR1* that confer caffeine resistance (Reinke et al. 2006; Moresi et al. 2023; Geck et al. 2024). Overexpression of the ABC transporters Snq2 and Pdr5 (Tsujimoto et al. 2015) and mutations in *PDR5* and *PDR1* have been found to increase yeast caffeine resistance (Sürmeli et al. 2019; Moresi et al. 2023; Geck et al. 2024), which supports our finding that the 12Δ strain, which lacks Pdr1, Pdr5, and Snq2, is sensitized to caffeine as well as the analog aminophylline. The Tor1-I1954V mutation previously identified to confer resistance to caffeine (Reinke et al. 2006) also confers resistance to caffeine in the sensitized 12Δ background. This provides support that similar mechanisms of resistance are retained in this strain background, which helps to support the use of this genetic background for drug mechanism of action discovery (Chinen et al. 2018). Despite the structural similarity of aminophylline to caffeine, the Tor1-I1954V mutation did not confer similar resistance to aminophylline. This suggests that the methyl group present on caffeine at N7, which is absent from aminophylline, may bind Tor1 within the FRB region containing I1954.

Caffeine extends the lifespan of yeast (Wanke et al. 2008; Rallis et al. 2013), and worms (Lublin et al. 2011; Sutphin et al. 2012; Bridi et al. 2015; Du et al. 2018; Li et al. 2019). Li et al. find that the lifespan effect of caffeine is dependent on the IGF-1 pathway (Du et al. 2018), which can act upstream of mTOR signaling (Johnson 2018). It remains of interest whether caffeine’s impact on longevity in these model organisms is directly or indirectly dependent on mTOR. *S. cerevisiae* Tor1-I1954V confers resistance to caffeine mediated growth inhibition (Reinke et al. 2006), and *C. elegans* contain a valine at the equivalent position in *let-363*, suggesting that the *C. elegans* mTOR protein may be more resistant to caffeine compared to yeast and humans. Mutation of this residue in *C. elegans* to alanine could be used to determine if it reduces the concentration of caffeine necessary for lifespan extension, which could provide evidence in regards to the role of the TOR pathway in caffeine mediated longevity in worms.

Aminophylline consists of the caffeine analog theophylline with ethylenediamine to improve solubility. Theophylline has been reported to extend worm lifespan (Li et al. 2019). Theophylline at 5 mM inhibits immunopurified mTOR in vitro and inhibits mTOR kinase activity in 3T3-L1 adipocyte cells (Scott and Lawrence 1998). Caffeine and theophylline are reported to inhibit mTOR as well as PI3K, with subunit p110δ most potently with an IC_50_ of 75µM (Foukas et al. 2002). Aminophylline has also been reported to reduce the elevated phospho-S2448 mTOR in an in vivo rat model of chronic renal failure (Liao et al. 2024).

In mammals, the primary molecular mechanisms of caffeine include phosphodiesterase inhibition, antagonism of adenosine receptors, and alterations in intracellular calcium (Daly 2000). Caffeine’s effects on PI3K may also be physiologically relevant (Foukas et al. 2002). Effects of caffeine on mTOR have been observed primarily at high concentrations, so the physiological relevance of mTOR inhibition in mammalian cells is unclear. Multiple beneficial effects have been observed for coffee consumption and human age-associated diseases as well as a reduction of all-cause mortality, however, many beneficial effects have also been observed with decaffeinated coffee (Van Dam et al. 2020), suggesting caffeine independent effects. This is further complicated as decaffeinated coffee still contains caffeine at lower concentrations (McCusker et al. 2006).

We also tested a variety of additional compounds for potential impacts on the TOR pathway using our yeast drug sensitive cell model. Canagliflozin has been found to impact the mTOR pathway (Jiang et al. 2023), and to extend lifespan of male mice (Miller et al. 2020, 2024). Nebivolol has been suggested to inhibit TORC1, but no lifespan extension was observed in mice under the conditions tested (Miller et al. 2024). Alpha lipoic acid may reduce mTOR signaling (Xie et al. 2012; Li et al. 2014). Isoliquiritigenin and withaferin A were also tested based on their predicted potential as rapamycin mimetic molecules (Aliper et al. 2017).

Ganoderic acid A was also tested as it is commercially available, and butylated ganoderic acid A was predicted to be an FKBP12-dependent mTOR inhibitor using a machine learning approach (Posansee et al. 2023).Taurine declines with age and supplementation can promote longevity in worms and mice, although the molecular mechanism is unclear (Singh et al. 2023). At the concentrations tested, taurine, ganoderic acid A, and alpha-lipoic acid had no effect on the 12Δ yeast strains, suggesting that the compounds may not inhibit yeast TOR and/or were unable to efficiently enter into yeast cells. Canagliflozin, isoliquiritigenin, withaferin A, and nebivolol at the concentrations tested had growth inhibitory effects in the 12Δ strain, however no observations of selective sensitivity of the *tor1* strain was observed. This suggests that the growth inhibitory activity of these compounds may be *TOR1*-independent. Canagliflozin, isoliquiritigenin, withaferin A, and nebivolol caused growth inhibition in 12Δ, but not the wildtype strain background, suggesting that they may be actively effluxed from yeast cells by ABC transporters that are absent in the 12Δ strain.

Using this yeast-based system for identifying *tor1*-dependent inhibitors requires an optimized concentration (*i*.*e*., range finding for optimal doses) for detection of the preferential inhibition of the *tor1* deficient strain. However, identifying *fpr1*-dependent inhibition such as that observed with rapamycin is observable over a much larger concentration range and can be more easily qualitatively identified. In addition to rapamycin, tacrolimus (FK506) is another natural product from *Streptomyces hygroscopicus* that inhibits growth in a FKBP12 dependent manner, yet the downstream molecular target of the drug-FKBP12 complex, calcineurin, is distinct. Of note, WDB0002, CEP250, meridamycin, and antascomycins A-E are additional FKBP12-dependent natural products that lack identified downstream targets (Rivera and Heitman 2023). Considering the robust rapamycin and rapalog resistance of the yeast *fpr1* mutant and the fact that various natural products can inhibit specific downstream targets through Fpr1-dependent mechanisms, it may be worthwhile to search for additional natural products with Fpr1-dependent effects on yeast growth.

Given the high-throughput capability of this system, molecular libraries including natural product libraries could be assayed using this approach. Future use of this system could also include identifying the effects of mixtures of compounds including assessing for synergistic interactions using yeast growth. Additionally, disrupting cell wall integrity (Chinen et al. 2014) could enhance the entry of compounds and further increase the sensitivity of the system.

## Methods

### Reagents

Reagents used in this study include rapamycin (LC Laboratories, Woburn, MA, USA), everolimus, temsirolimus, ridaforolimus, AZD8055, Torin1, GSK, canagliflozin hemihydrate, alpha lipoic acid, nebivolol (APExBIO, Houston, TX, USA), caffeine, theophylline, taurine (Thermo Fisher Scientific, Ward Hill, MA, USA), withaferin A (Adipogen, San Diego, CA, USA), ganoderic acid A (MCE, Monmouth Junction, NJ, USA), isoliquiritigenin, and aminophylline (TCI, Tokyo, Japan).

### Yeast strains

Yeast strains used in this study are shown in Supplemental Table 2. CRISPR was used for yeast strain generation similarly to as previously described (Laughery et al. 2015; Laughery and Wyrick 2019; Shortt et al. 2023). Briefly, sgRNA targeting sequences were selected using Geneious Prime software, and were engineered to contain SmiI and BclI sites for cloning. Oligonucleotides (Azenta Genewiz, South Plainfield, NJ, USA) containing the sgRNA targeting sequence were hybridized to double-stranded DNA and then ligated using T4 DNA ligase into SmiI and BclI (NEB, Ipswich, MA, USA) digested pML104. Plasmids were confirmed to contain the desired sgRNA by PCR verification. Oligonucleotide sequences used in this study are shown in Supplemental Table 3. Yeast strains (either BY4742 or the 12Δ strain) were transformed using a EZ Yeast Transformation kit (Zymo Research Corporation, Irvine, CA, USA) with the indicated sgRNA targeting CRISPR plasmid along with a corresponding repair template. Sanger sequencing was performed to verify the presence of the desired mutations. The Saccharomyces Genome Database (SGD) was a critical resource used in support of this study (Wong et al. 2023).

### Yeast liquid culture growth assays

Overnight cultures were inoculated using a single colony and grown at 30 °C with shaking in YPD media consisting of 1% Bacto yeast extract (BD, Franklin Lakes, NJ, USA), 2% Bacto peptone (BD, Franklin Lakes, NJ, USA), and 2% dextrose (Sunrise Science Products, Knoxville, TN, USA). The following day, the overnight culture was diluted ∼1/100 into YPD and all samples were adjusted to an equal initial optical density (OD) of 0.10 +/-0.02 using a Biochrom WPA Biowave CO8000 cell density meter. A 150 μL aliquot of the diluted overnight culture was added to 96 well plates, with the exception of blanks, which contained media only. To the samples, a 1.5 μL aliquot of respective chemical compound was added to each treatment group with the exception of caffeine and aminophylline, of which 15 μL of drug was added.

DMSO vehicle controls were included where appropriate. Growth assays were performed using an Epoch2 Microplate Reader (BioTek, Winooski, VT, USA) with Gen6 software version 1.03.01. 96-well microplates were incubated at 30 °C with continuous double orbital shaking with a frequency of 807 cycles per minute (cpm). A lid temperature gradient was set to + 2 °C to prevent condensation, and absorbance at 600 nm (OD600) was measured every 30 mins for 24 h. Of note, 384-well plates were also tested with settings as above except with 60 µl volume and 269 cpm (6mm orbital) shaking based on a previous report (Jung et al. 2015), which was found to generate fermentative growth curves in the BY4742 strain backgrounds (data not shown). A standardized blank OD value was determined by taking the mean value of all blanks across all experiments and subtracting from all datasets. Dose response curves were first performed to identify the concentrations used within this study. All individual experiments were performed using replicates across 3 wells which were averaged, and each experiment was repeated independently three or more times. Individual representative experimental data are shown.

## Acknowledgments

The 12geneΔ0hsr strain (referred to within this manuscript as “12Δ”) was graciously provided by Dr. Takeo Usai from the University of Tsukuba. Funding for this project was provided by Longevity Impetus Grants (Norn Group) and an intramural grant from the Western University of Health Sciences. The authors have no conflicts of interest to declare.

## References

Aliper A, Jellen L, Cortese F, et al (2017) Towards natural mimetics of metformin and rapamycin. Aging (Albany NY) 9:2245–2268. 10.18632/aging.101319

Arriola Apelo SI, Neuman JC, Baar EL, et al (2016) Alternative rapamycin treatment regimens mitigate the impact of rapamycin on glucose homeostasis and the immune system. Aging Cell 15:28–38. 10.1111/acel.12405

Baghdadi M, Nespital T, Monzó C, et al (2024) Intermittent rapamycin feeding recapitulates some effects of continuous treatment while maintaining lifespan extension. Molecular Metabolism 81:101902. 10.1016/j.molmet.2024.101902

Biomedical O (2024) Million Molecule Challenge Results and Leaderboard – Ora Biomedical, Inc. In: Million Molecule Challenge Results and Leaderboard – Ora Biomedical, Inc. https://orabiomedical.com/mmcleaderboard/. 12 Jul 2024

Bitto A, Ito TK, Pineda VV, et al (2016) Transient rapamycin treatment can increase lifespan and healthspan in middle-aged mice. Elife 5:e16351. 10.7554/eLife.16351

Bjedov I, Toivonen JM, Kerr F, et al (2010) Mechanisms of Life Span Extension by Rapamycin in the Fruit Fly Drosophila melanogaster. Cell Metabolism 11:35–46. 10.1016/j.cmet.2009.11.010

Blagosklonny MV (2023) Cancer prevention with rapamycin. Oncotarget 14:342–350. 10.18632/oncotarget.28410

Bridi JC, Barros AG de A, Sampaio LR, et al (2015) Lifespan Extension Induced by Caffeine in Caenorhabditis elegans is Partially Dependent on Adenosine Signaling. Frontiers in Aging Neuroscience 7:

Chinen T, Hamada K, Taguchi A, et al (2018) Multidrug Sensitive Yeast Strains, Useful Tools for Chemical Genetics. In: Abdulkhair WMH (ed) The Yeast Role in Medical Applications. InTech

Chinen T, Nagumo Y, Usui T (2014) Construction of a genetic analysis-available multidrug sensitive yeast strain by disruption of the drug efflux system and conditional repression of the membrane barrier system. J Gen Appl Microbiol 60:160–162. 10.2323/jgam.60.160

Chresta CM, Davies BR, Hickson I, et al (2010) AZD8055 Is a Potent, Selective, and Orally Bioavailable ATP-Competitive Mammalian Target of Rapamycin Kinase Inhibitor with In vitro and In vivo Antitumor Activity. Cancer Research 70:288–298. 10.1158/0008-5472.CAN-09-1751

Daly JW (2000) Alkylxanthines as research tools. Journal of the Autonomic Nervous System 81:44–52. 10.1016/S0165-1838(00)00110-7

Du X, Guan Y, Huang Q, et al (2018) Low Concentrations of Caffeine and Its Analogs Extend the Lifespan of Caenorhabditis elegans by Modulating IGF-1-Like Pathway. Front Aging Neurosci 10:211. 10.3389/fnagi.2018.00211

Foukas LC, Daniele N, Ktori C, et al (2002) Direct Effects of Caffeine and Theophylline on p110δ and Other Phosphoinositide 3-Kinases. Journal of Biological Chemistry 277:37124–37130. 10.1074/jbc.M202101200

Friedman DB, Johnson TE (1988) A mutation in the age-1 gene in Caenorhabditis elegans lengthens life and reduces hermaphrodite fertility. Genetics 118:75–86. 10.1093/genetics/118.1.75

Gaubitz C, Prouteau M, Kusmider B, Loewith R (2016) TORC2 Structure and Function. Trends in Biochemical Sciences 41:532–545. 10.1016/j.tibs.2016.04.001

Geck RC, Moresi NG, Anderson LM, et al (2024) Experimental evolution of S. cerevisiae for caffeine tolerance alters multidrug resistance and TOR signaling pathways. G3 (Bethesda) jkae148. 10.1093/g3journal/jkae148

Harrison DE, Strong R, Sharp ZD, et al (2009) Rapamycin fed late in life extends lifespan in genetically heterogeneous mice. Nature 460:392–395. 10.1038/nature08221

Heitman J, Movva NR, Hall MN (1991) Targets for cell cycle arrest by the immunosuppressant rapamycin in yeast. Science (New York, NY) 253:905–9

Jiang E, Dinesh A, Jadhav S, et al (2023) Canagliflozin shares common mTOR and MAPK signaling mechanisms with other lifespan extension treatments. Life Sci 328:121904. 10.1016/j.lfs.2023.121904

Johnson SC (2018) Nutrient Sensing, Signaling and Ageing: The Role of IGF-1 and mTOR in Ageing and Age-Related Disease. Subcell Biochem 90:49–97. 10.1007/978-981-13-2835-0_3

Jung PP, Christian N, Kay DP, et al (2015) Protocols and Programs for High-Throughput Growth and Aging Phenotyping in Yeast. PLOS ONE 10:e0119807. 10.1371/journal.pone.0119807

Kaeberlein M, Powers RW, Steffen KK, et al (2005) Regulation of yeast replicative life span by TOR and Sch9 in response to nutrients. Science 310:1193–1196. 10.1126/science.1115535

Kaeberlein TL, Green AS, Haddad G, et al (2023) Evaluation of off-label rapamycin use to promote healthspan in 333 adults. Geroscience 45:2757–2768. 10.1007/s11357-023-00818-1

Kapahi P, Zid BM, Harper T, et al (2004) Regulation of Lifespan in Drosophila by Modulation of Genes in the TOR Signaling Pathway. Current Biology 14:885–890. 10.1016/j.cub.2004.03.059

Knight SD, Adams ND, Burgess JL, et al (2010) Discovery of GSK2126458, a Highly Potent Inhibitor of PI3K and the Mammalian Target of Rapamycin. ACS Med Chem Lett 1:39–43. 10.1021/ml900028r

Lamming DW, Ye L, Katajisto P, et al (2012) Rapamycin-Induced Insulin Resistance Is Mediated by mTORC2 Loss and Uncoupled from Longevity. Science 335:1638–1643. 10.1126/science.1215135

Laughery MF, Hunter T, Brown A, et al (2015) New vectors for simple and streamlined CRISPR-Cas9 genome editing in Saccharomyces cerevisiae. Yeast 32:711–720. 10.1002/yea.3098

Laughery MF, Wyrick JJ (2019) Simple CRISPR-Cas9 Genome Editing in Saccharomyces cerevisiae. Curr Protoc Mol Biol. 10.1002/cpmb.110

Lee DJW, Kuerec AH, Maier AB (2024) Targeting ageing with rapamycin and its derivatives in humans: a systematic review. The Lancet Healthy Longevity 5:e152–e162. 10.1016/S2666-7568(23)00258-1

Lee MB, Carr DT, Kiflezghi MG, et al (2017) A system to identify inhibitors of mTOR signaling using high-resolution growth analysis in Saccharomyces cerevisiae. Geroscience 39:419–428. 10.1007/s11357-017-9988-4

Li H, Roxo M, Cheng X, et al (2019) Pro-oxidant and lifespan extension effects of caffeine and related methylxanthines in Caenorhabditis elegans. Food Chemistry: X 1:100005. 10.1016/j.fochx.2019.100005

Li Z, Dungan CM, Carrier B, et al (2014) Alpha-lipoic acid supplementation reduces mTORC1 signaling in skeletal muscle from high fat fed, obese Zucker rats. Lipids 49:1193–1201. 10.1007/s11745-014-3964-x

Liao X, Lu J, Huang Z, et al (2024) Aminophylline suppresses chronic renal failure progression by activating SIRT1/AMPK/mTOR-dependent autophagy. ABBS. 10.3724/abbs.2024049

Liu GY, Sabatini DM (2020) mTOR at the nexus of nutrition, growth, ageing and disease. Nat Rev Mol Cell Biol 21:183–203. 10.1038/s41580-019-0199-y

Liu Q, Chang JW, Wang J, et al (2010) Discovery of 1-(4-(4-Propionylpiperazin-1-yl)-3-(trifluoromethyl)phenyl)-9-(quinolin-3-yl)benzo[h][1,6]naphthyridin-2(1 H)-one as a Highly Potent, Selective Mammalian Target of Rapamycin (mTOR) Inhibitor for the Treatment of Cancer. J Med Chem 53:7146–7155. 10.1021/jm101144f

Lublin A, Isoda F, Patel H, et al (2011) FDA-approved drugs that protect mammalian neurons from glucose toxicity slow aging dependent on cbp and protect against proteotoxicity. PLoS One 6:e27762. 10.1371/journal.pone.0027762

Lucanic M, Plummer WT, Chen E, et al (2017) Impact of genetic background and experimental reproducibility on identifying chemical compounds with robust longevity effects. Nat Commun 8:14256. 10.1038/ncomms14256

McCusker RR, Fuehrlein B, Goldberger BA, et al (2006) Caffeine Content of Decaffeinated Coffee. Journal of Analytical Toxicology 30:611–613. 10.1093/jat/30.8.611

Miller RA, Harrison DE, Allison DB, et al (2020) Canagliflozin extends life span in genetically heterogeneous male but not female mice. JCI Insight 5:e140019. 10.1172/jci.insight.140019

Miller RA, Harrison DE, Cortopassi GA, et al (2024) Lifespan effects in male UM-HET3 mice treated with sodium thiosulfate, 16-hydroxyestriol, and late-start canagliflozin. GeroScience. 10.1007/s11357-024-01176-2

Moresi NG, Geck RC, Skophammer R, et al (2023) Caffeine-tolerant mutations selected through an at-home yeast experimental evolution teaching lab. 10.1101/2023.01.17.524437

Morris JZ, Tissenbaum HA, Ruvkun G (1996) A phosphatidylinositol-3-OH kinase family member regulating longevity and diapause in Caenorhabditis elegans. Nature 382:536–539. 10.1038/382536a0

Phelps GB, Morin J, Pinto C, et al (2024) Comprehensive evaluation of lifespan-extending molecules in C. elegans. 2024.06.24.600458

Phillips EJ, Simons MJP (2023) Rapamycin not dietary restriction improves resilience against pathogens: a meta-analysis. Geroscience 45:1263–1270. 10.1007/s11357-022-00691-4

Pitt JN, Strait NL, Vayndorf EM, et al (2019) WormBot, an open-source robotics platform for survival and behavior analysis in C. elegans. GeroScience 41:961–973. 10.1007/s11357-019-00124-9

Popova NV, Jücker M (2021) The Role of mTOR Signaling as a Therapeutic Target in Cancer. International Journal of Molecular Sciences 22:1743. 10.3390/ijms22041743

Posansee K, Liangruksa M, Termsaithong T, et al (2023) Combined Deep Learning and Molecular Modeling Techniques on the Virtual Screening of New mTOR Inhibitors from the Thai Mushroom Database. ACS Omega 8:38373–38385. 10.1021/acsomega.3c04827

Powers RW, Kaeberlein M, Caldwell SD, et al (2006) Extension of chronological life span in yeast by decreased TOR pathway signaling. Genes & Development 20:174–184. 10.1101/gad.1381406

Rallis C, Codlin S, Bähler J (2013) TORC1 signaling inhibition by rapamycin and caffeine affect lifespan, global gene expression, and cell proliferation of fission yeast. Aging Cell 12:563–573. 10.1111/acel.12080

Reinke A, Chen JC-Y, Aronova S, Powers T (2006) Caffeine Targets TOR Complex I and Provides Evidence for a Regulatory Link between the FRB and Kinase Domains of Tor1p. Journal of Biological Chemistry 281:31616–31626. 10.1016/S0021-9258(19)84075-9

Rivera A, Heitman J (2023) Natural product ligands of FKBP12: Immunosuppressive antifungal agents FK506, rapamycin, and beyond. PLoS Pathog 19:e1011056. 10.1371/journal.ppat.1011056

Sarbassov DD, Ali SM, Sengupta S, et al (2006) Prolonged rapamycin treatment inhibits mTORC2 assembly and Akt/PKB. Mol Cell 22:159–168. 10.1016/j.molcel.2006.03.029

Scott PH, Lawrence JC (1998) Attenuation of Mammalian Target of Rapamycin Activity by Increased cAMP in 3T3-L1 Adipocytes. Journal of Biological Chemistry 273:34496–34501. 10.1074/jbc.273.51.34496

Shortt C, Krippaehne E, Wasko BM (2023) A simple and accessible CRISPR genome editing laboratory exercise using yeast. MicroPubl Biol 2023:. 10.17912/micropub.biology.000699

Singh P, Gollapalli K, Mangiola S, et al (2023) Taurine deficiency as a driver of aging. Science 380:eabn9257. 10.1126/science.abn9257

Sürmeli Y, Holyavkin C, Topaloğlu A, et al (2019) Evolutionary engineering and molecular characterization of a caffeine-resistant Saccharomyces cerevisiae strain. World J Microbiol Biotechnol 35:183. 10.1007/s11274-019-2762-2

Sutphin GL, Bishop E, Yanos ME, et al (2012) Caffeine extends life span, improves healthspan, and delays age-associated pathology in Caenorhabditis elegans. Longevity & Healthspan 1:9. 10.1186/2046-2395-1-9

Tsujimoto Y, Shimizu Y, Otake K, et al (2015) Multidrug resistance transporters Snq2p and Pdr5p mediate caffeine efflux in Saccharomyces cerevisiae. Bioscience, Biotechnology, and Biochemistry 79:1103–1110. 10.1080/09168451.2015.1010476

Van Dam RM, Hu FB, Willett WC (2020) Coffee, Caffeine, and Health. N Engl J Med 383:369–378. 10.1056/NEJMra1816604

Vellai T, Takacs-Vellai K, Zhang Y, et al (2003) Influence of TOR kinase on lifespan in C. elegans. Nature 426:620–620. 10.1038/426620a

Wanke V, Cameroni E, Uotila A, et al (2008) Caffeine extends yeast lifespan by targeting TORC1. Molecular Microbiology 69:277–285. 10.1111/j.1365-2958.2008.06292.x

Wong ED, Miyasato SR, Aleksander S, et al (2023) Saccharomyces genome database update: server architecture, pan-genome nomenclature, and external resources. Genetics 224:iyac191. 10.1093/genetics/iyac191

Wu JJ, Liu J, Chen EB, et al (2013) Increased Mammalian Lifespan and a Segmental and Tissue-Specific Slowing of Aging after Genetic Reduction of mTOR Expression. Cell Reports 4:913–920. 10.1016/j.celrep.2013.07.030

Xie R, Li X, Ling Y, et al (2012) Alpha-lipoic acid pre- and post-treatments provide protection against in vitro ischemia-reperfusion injury in cerebral endothelial cells via Akt/mTOR signaling. Brain Research 1482:81–90. 10.1016/j.brainres.2012.09.009

