## Supplemental Data for "An mTOR Inhibitor Discovery System Using the Growth of Drug-Sensitized Yeast Strains"

**Supplemental Table 1. Effects of Select Compounds on Yeast Growth**

| Compound Name | Maximum Concentration Tested | Strain Background Tested | Impact on Yeast Growth |
|---|---|---|---|
| Ganoderic Acid A | 20 µM | 12D | no growth inhibition |
| α-Lipoic Acid | 100 µM | 12d | no growth inhibition |
| Taurine | 50 mM | 12d | no growth inhibition |
| Canagliflozin and Canagliflozin Hemihydrate | 100 µM | 12d | growth inhibition, but no tor1 sensitivity |
| | 100 µM | WT | no growth inhibition |
| Nebivolol | 25 µM | 12d | growth inhibition, but no tor1 sensitivity |
| | 25 µM | WT | no growth inhibition |
| Isoliquiritigenin | 780 µM | 12d | growth inhibition, but no tor1 sensitivity |
| | 780 µM | WT | no growth inhibition |
| Withaferin A | 20 µM | 12d | growth inhibition, but no tor1 sensitivity |
| | 20 µM | WT | no growth inhibition |

**Supplemental Table 2. Yeast strains used in this study.**

| Yeast strain | Additional identifier | Genotype | protein change | Reference/Source |
|---|---|---|---|---|
| BY4742 | BY4742 | MATα *ura3Δ his3Δ1 leu2Δ lys2Δ* | | Brachmann et al. |
| BW746 | *tor1* | MATa, *tor1Δ::URA3 his3Δ1 leu2Δ* | | Kaeberlein lab |
| BW1381 | *fpr1* | MATα *fpr1Δ::KanMX* | | Lee et al. |
| BW1443 | *tor1* | as BY4742, *tor1-G64T,C66A* | Tor1-D22* | this study |
| BW1444 | *tor1-1* | as BY4742, *tor1-C5916A* | Tor1-S1972R | this study |
| BW1440 | Tor1-I1954V | as BY4742, *tor1-C5832T,A5860G* | Tor1-I1954V | this study |
| OTA017 (BW1457) | 12Δ (12geneΔ0HSR) | *MATa his3Δ1 leu2Δ0 met15Δ0 ura3Δ0 pdr3Δ0 pdr8Δ0 pdr1Δ0 yrr1Δ0 snq2Δ0 pdr5Δ0 pdr10Δ0 yor1Δ0 pdr15Δ0 pdr11Δ0 pdr12Δ0 aus1Δ0 RME1(ins308A)* | | Chinen et al. |
| BW1458 | 12Δ *tor1* | as 12Δ , *tor1-G64T,C66A* | Tor1-D22* | this study |
| BW1460 | 12Δ *tor1-1* | as 12Δ , t*or1-C5916A* | Tor1-S1972R | this study |
| BW1462 | 12Δ *fpr1* | as 12Δ , *fpr1-G55T,T57A* | Fpr1-G19* | this study |
| BW1463 | 12Δ Tor1-I1954V | as 12Δ , *tor1-C5832T,A5860G* | Tor1-I1954V | this study |

References:

Brachmann CB et al. *Yeast*. 1998 Jan 30;14(2):115-32. PMID9483801
Chinen T et al. *J Gen Appl Microbiol*. 2014;60(4):160-2. PMID25273990
Lee MB et al. *Geroscience. 2017* Aug;39(4):419-428. PMID28707282

Supplemental Table 3. Oligonucelotides and plasmids used in this study for yeast strain generation.

| Oligo | Oligonucleotide Sequence |
|---|---|
| oBW734 | GATCCGTGACTAACTAATTCTGCCGTTTTAGAGCTAG |
| oBW735 | CTAGCTCTAAAACGGCAGAATTAGTTAGTCACG |
| oBW736 | GATTCATAGTCCAGTCCTGGTAAATCAGGCAGAATTAGTTAGTCACGAGTTGGTCAGAGTAGCCGTTCTATGGCACG |
| oBW737 | CGTGCCATAGAACGGCTACTCTGACCAACTCGTGACTAACTAATTCTGCCTGATTTACCAGGACTGGACTATGAATC |
| oBW742 | GATCGAAAGCGGCTAACAACGATAGTTTTAGAGCTAG |
| oBW743 | CTAGCTCTAAAACTATCGTTGTTAGCCGCTTTC |
| oBW744 | AACTTTTGAAAGCGGCTAACAACGATATGTAAATGGATAGAAATGTGCCGTTGGCACCGAAT |
| oBW745 | ATTCGGTGCCAACGGCACATTTCTATCCATTTACATATCGTTGTTAGCCGCTTTCAAAAGTT |
| oBW746 | GATCTATGTTCAACGAAAAATTGGGTTTTAGAGCTAG |
| oBW747 | CTAGCTCTAAAACCCAATTTTTCGTTGAACATA |
| oBW748 | TTATGGTATGAAGGACTGGAAGATGCGAGACGCCAATTTTTCGTTGAACATAACATAGAA |
| oBW749 | TTCTATGTTATGTTCAACGAAAAATTGGCGTCTCGCATCTTCCAGTCCTTCATACCATAA |
| oBW822 | GATCGACAGAATTTCCCCAGGTGAGTTTTAGAGCTAG |
| oBW823 | CTAGCTCTAAAACTCACCTGGGGAAATTCTGTC |
| oBW824 | CAAAATTGACAGAATTTCCCCAGGTGATTGAGCCACCTTCCCAAAGACAGGTGACTTGGTTA |
| oBW825 | TAACCAAGTCACCTGTCTTTGGGAAGGTGGCTCAATCACCTGGGGAAATTCTGTCAATTTTG |

| Plasmid | sgRNA target site on gDNA | Plasmid cloning oligos | Plasmid backbone |
|---|---|---|---|
| pBW280 | CGTGACTAACTAATTCTGCCTGG | oBW734+oBW735 | pML104 |
| pBW281 | GAAAGCGGCTAACAACGATATGG | oBW742+oBW743 | pML104 |
| pBW282 | TATGTTCAACGAAAAATTGGCGG | oBW746+oBW747 | pML104 |
| pBW292 | GACAGAATTTCCCCAGGTGATGG | oBW822+oBW823 | pML104 |

| Yeast Strain | Plasmid used to make strain | Repair Template Oligos used to make strain |
|---|---|---|
| BW1440 | pBW280 | oBW736 & oBW737 |
| BW1463 | pBW280 | oBW736 & oBW737 |
| BW1443 | pBW281 | oBW744 & oBW745 |
| BW1458 | pBW281 | oBW744 & oBW745 |
| BW1444 | pBW282 | oBW748 & oBW749 |
| BW1460 | pBW282 | oBW748 & oBW749 |
| BW1462 | pBW292 | oBW824 & oBW825 |
